# Anti-Quorum Sensing Phages Disarm *Pseudomonas aeruginosa*

**DOI:** 10.64898/2026.02.16.706229

**Authors:** Jesper Juel Mauritzen, Kira Céline Koonce, Swenja Lassen, Nina Molin Høyland-Kroghsbo

**Author notes:** These authors contributed equally (Jesper Juel Mauritzen, Kira Céline Koonce). Address correspondence to Nina Molin Høyland-Kroghsbo.

## Abstract

By 2050, the death toll from previously preventable or easily curable bacterial infections is projected to surpass that caused by cancer, unless we prevent the spread of antibiotic resistance and develop new therapies. A promising approach is phage therapy, which exploits bacteriophages, natural predators of bacteria. However, bacteria fight back, which can limit its efficacy. Notably, many bacteria rely on cell-cell communication, known as quorum sensing, to orchestrate both virulence programs and phage defenses. To circumvent these, we have engineered anti-quorum sensing phages against the human pathogen *Pseudomonas aeruginosa*. Our engineered phages effectively degrade quorum-sensing molecules, reduce virulence factor production, and double the survival of *P. aeruginosa*-infected *Galleria mellonella* larvae. Moreover, we demonstrate that the anti-quorum sensing phages inhibit quorum sensing in mixed populations of phage-susceptible and phage-resistant cells, demonstrating the ability of the phages to disarm subpopulations phage-resistant *P. aeruginosa*, which often are selected for during phage treatment. Together, our findings highlight the future therapeutic promise of anti-quorum sensing phages as a dual-action strategy in killing susceptible cells while attenuating virulence across the bacterial population. This approach has the potential to enhance the robustness of phage therapy.

## Introduction

The alarming rise of antibiotic resistance calls for an urgent need to develop alternative therapies. The increasingly antibiotic-resistant pathogenic bacterium *Pseudomonas aeruginosa* is one of the “priority pathogens” that pose the greatest threat to human health [1]. *P. aeruginosa* causes severe disease in chronic cystic fibrosis lung infections and is a major source of hospital acquired infections in *e.g*. burn wound victims and immunocompromised patients [2, 3]. Moreover, *P. aeruginosa* is a versatile pathogen, infecting animals and plants, underscoring its importance for agriculture and food security [4].

The use of bacteriophage viruses (phages) as an option for killing pathogenic bacteria is reemerging. Phages are viruses that infect, replicate within, and ultimately kill bacteria. During a single lytic infectious cycle, a phage may produce about 100-150 phage progeny, which are released when the bacterium lyses. Alternatively, temperate phages can execute the lysogenic life cycle, integrating their genome into the host chromosome to form a lysogen, where they reside dormant as a prophage until cellular or environmental conditions trigger a switch to lytic replication. Shortly after the discovery of phages over 100 years ago, phage therapy was applied clinically to treat bacterial infections. However, limited understanding of phage-host interactions and thus inconsistent therapeutic outcomes meant that phage therapy had limited success and was largely abandoned in the wake of antibiotics [5], except at key phage therapy institutes such as the George Eliava Institute of Bacteriophages [6]. In the context of the escalating and global threat of antibiotic resistance, phage therapy is now re-emerging as a promising alternative. In recent years, phage therapies using natural virulent phages or phages engineered to be exclusively lytic, have been used for compassionate care use, successfully having treated several critically ill patients suffering from multidrug-resistant infections. However, in some cases, targeted bacteria developed resistance to the administrated phages [7–9]. Although cocktails composed of multiple phages that target the same pathogen via different phage receptors reduce the probability that bacteria simultaneously evolve resistance to those phages, multi-phage-resistant bacteria can still emerge [10].

In the evolutionary arms race with phages, bacteria have developed a range of defense mechanisms. These include so-called primary defenses, such as phage receptor modifications or loss, which prevent phages from infecting the bacteria in the first place. If these defenses fail, bacteria deploy secondary defense systems intracellularly; blocking of phage replication or preventing the phage from converting the bacterial cell into a viral factory [11]. Intracellular phage defenses are diverse and plentiful [12] and include the Clustered Regularly Interspaced Short Palindromic Repeats (CRISPR) system with its CRISPR-Associated (Cas) proteins; microbial adaptive immune systems [13]. While CRISPR-Cas function at the individual cell level, bacterial defenses are not limited to that.

In response to the employment of CRISPR-Cas immunity, phages have evolved mechanisms of counter defense, neutralizing bacterial adaptive immunity. These include anti-CRISPR (Acr) proteins [14] which inhibit CRISPR-Cas through diverse mechanisms, such as blocking DNA binding, preventing target cleavage, and interfering with Cas complex assembly [15]. Acr genes are typically encoded in operons driven by a strong acr-associated promoter (P_acr_), enabling rapid, high-level *acr* transcription immediately after phage DNA enters the cell. This rapid accumulation of Acr proteins ultimately allows overcoming host CRISPR defense [16]. While anti-CRISPR proteins provide a powerful tool of phages to effectively neutralize host immunity, their expression must be tightly controlled as unrestrained Acr production can be lethal to the phage. Thus, to prevent overexpression, acr operons additionally encode an anti-CRISPR-associated (aca) gene directly repressing P_acr_-mediated transcription.

Similarly, the inappropriate expression of host defense systems is costly. Thus, bacteria have developed mechanisms to regulate their defenses in accordance with the risk of infection. For example, quorum sensing (QS), a cell-cell communication mechanism, enables bacteria to monitor cell density and behave as if they were a multicellular organism. QS depends on the production, release, and group-wide detection of diffusible QS signaling molecules. At high-cell-density, the accumulation of AIs induces bacterial populations to synchronously activate hundreds of genes involved in group behavior, such as virulence genes and those encoding phage defenses [17–22].

*P. aeruginosa* communicates using QS to coordinate behavior such as biofilm formation and virulence in addition to launching anti-phage defense systems [18, 20, 23]. As described above, anti-phage defenses [12], some of which are QS-regulated [18–22, 24], pose one of the major biological challenges to phage therapy and biocontrol. *P. aeruginosa* has multiple intertwined QS systems, including two canonical LuxI/R-type QS systems: LasI/R and RhlI/R. LasI/R is conventionally considered at the top of the QS cascade in *P. aeruginosa*. Laslsynthesizes 3-Oxo-C12-homoserine lactone (3O-C12-HSL), which is detected by LasR. 3OC12-HSL-activated LasR turns on transcription of the QS circuit genes *rhlI/R* and *pqsABCDE, pqsH*, and *pqsR*. RhlI/R, in turn, produces and detects C4-homoserine lactone (C4-HSL). The Pseudomonas quinolone signal (PQS) system produces and detects 2-heptyl-4-quinolone (HHQ) and 2-heptyl-3,4-dihydroxyquinoline (PQS) [25–28]. Collectively, these QS systems control the production of an array of virulence factors including elastase, rhamnolipids, and the green pigment pyocyanin [23]. Intriguingly, while the Las QS system generally activates the PQS system, phage infection of LasI deficient mutants bypasses this regulatory link and directly activates PQS QS [29]. PQS produced by phage-infected cells, allows healthy *P. aeruginosa* swarms to avoid colliding with phage-infected cells[18]. Thus, phage infection directly activates PQS QS, which controls virulence factors and phage-defense systems, which may limit successful outcomes of phage therapy.

QS inhibitors indeed block bacterial virulence of various pathogens [30] and downregulate the CRISPR-Cas phage defense [20], thus muting bacteria may simultaneously disarm bacterial virulence programs and anti-phage defenses. Here, we combine QS inhibition and phage therapy directly by engineering phages to deliver enzymes that degrade homoserine lactones and PQS in *P. aeruginosa* [31].

## Results

*P. aeruginosa* uses QS to coordinate behavior such as biofilm formation and virulence in addition to launching anti-phage defense systems [18, 20, 23]. Thus, inhibiting QS can disarm the pathogen without killing it, offering an effective anti-virulence strategy [30]. Another promising antimicrobial approach is phage therapy, which directly kills bacteria [9]. Here, we combined these two treatment strategies by engineering the *P. aeruginosa* phage DMS3 (accession no. NC_008717) to express the N-acyl homoserine lactonase QsdA, which cleaves AHLs [32], and the PQS dioxygenase AqdC, which cleaves PQS [33], and reduces *P. aeruginosa* virulence [34]. To ensure potent *qsdA* and *aqdC* expression, we cloned the genes under the DMS3 P_*acr*_ promoter, replacing the native anti-CRISPR gene *acrIE*. This promoter drives strong early expression upon phage infection, but its activity is later repressed by the Aca repressor [16]. The DMS3 phage naturally encodes the anti-QS peptide Aqs1 that binds and inhibits LasR [35]. To isolate and assess the anti-QS effects of expressing QS-degrading enzymes from DMS3, we removed this native LasR inhibiting activity by mutating the *aqs1* gene, preventing Aqs1 from binding and blocking LasR [35]. Finally, we mutated the *cI* repressor in DMS3 to force the phage into a fully lytic lifecycle [36].

### Anti-QS phages inhibit C4-HSL and PQS accumulation

First, we tested our engineered DMS3 phages; DMS3 *aqs1_FSDARE* vir (DMS3 parent hereafter), DMS3 *aqs1_FSDARE* Δ*acrIE*::*qsdA* vir (DMS3 QsdA hereafter) and DMS3 *aqs1_FSDARE* Δ*acrIE*::*aqdC* vir (DMS3 AqdC hereafter); for their ability to inhibit QS molecule accumulation. To do this, we cultured *P. aeruginosa* UCBPP-PA14 (PA14 hereafter) to early exponential phase and either left them uninfected or challenged with the parental DMS3, DMS3 QsdA, AqdC or a 1:1 mixture of both anti-QS phages at an MOI of 10 (Fig. 1). Extracellular 3OC12-HSL, C4-HSL, and PQS were quantified in sterile-filtered supernatants using *Escherichia coli* bioreporters specific to each molecule. Each bioreporter strain carried either a *lasR, rhlR*, or *pqsR* pBAD expression vector and a *lasB, rhlA*, or *pqsA* promoter, respectively, upstream of the *luxCDABE* operon in pCS26 [37, 38].

**Figure 1.**
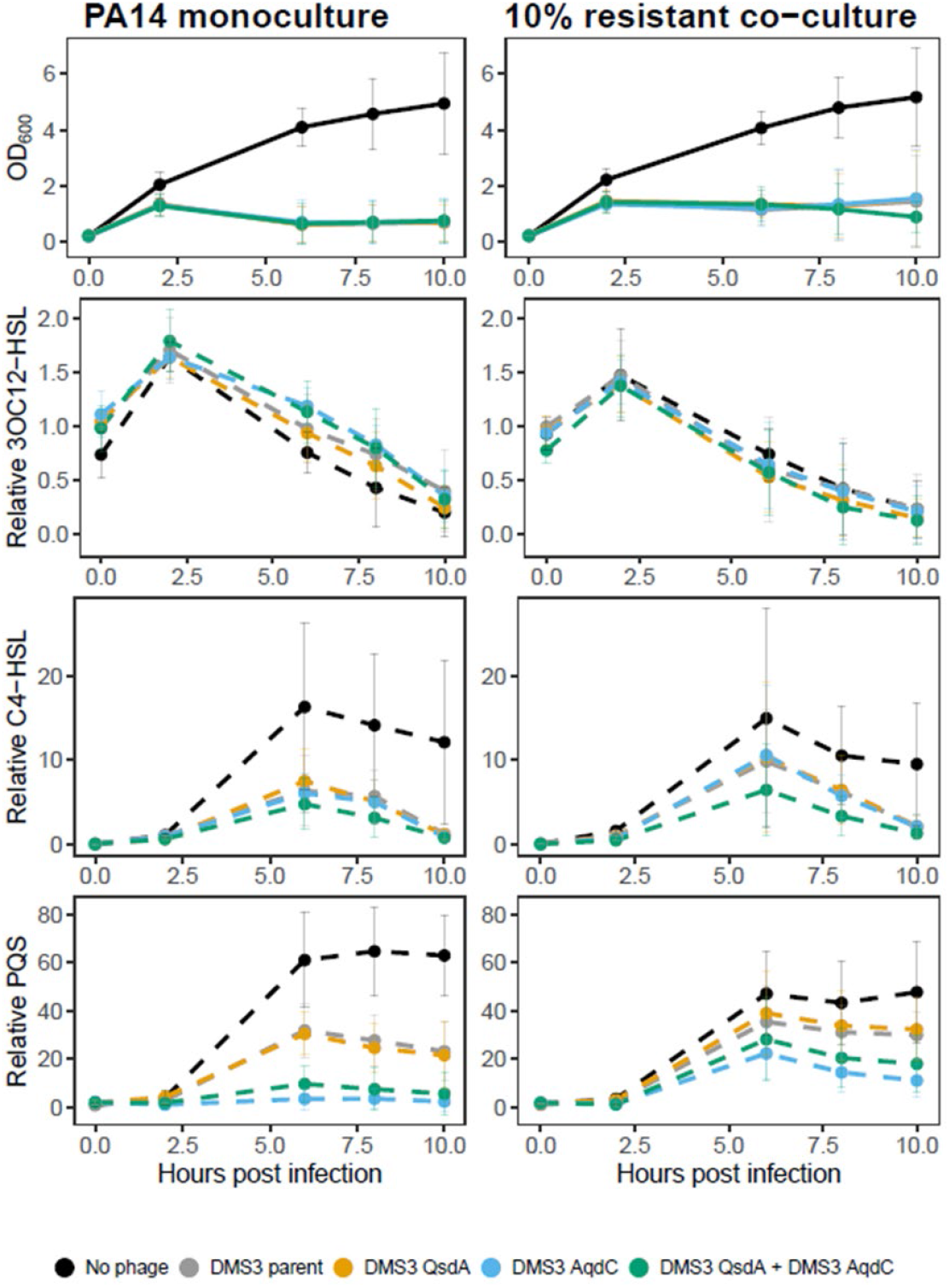
Anti-QS phages reduce C4-HSL and PQS accumulation. PA14 monocultures and 10% resistant co-cultures (90% PA14; 10% PA14^R^) were infected at OD_600_ ~ 0.2 with parental DMS3 phage or anti-QS phages expressing *qsdA, aqdC* or a 1:1 mixture of both anti-QS phages (MOI=10; 2*10^9^ PFU ml^-1^). Bacterial abundance was monitored by OD_600_, and relative QS molecule levels were quantified using *E. coli* bioreporters. Error bars represent standard deviation from biological replicates (*n*=4). Pairwise treatment differences were estimated using generalized additive mixed models (Figure S1).

The anti-QS phages killed PA14 to a similar extent as the parental DMS3 phage, indicating that replacing the native anti-CRISPR gene *acrIE* with *qsdA* or *aqdC* genes did not disrupt phage replication nor improve bacterial killing (Fig. 1). While infection with anti-QS phages did not significantly reduce 3OC12-HSL accumulation under the experimental conditions (Fig. 1 and S1), the combination of DMS3 QsdA and AqdC significantly lowered C4-HSL levels, reducing accumulation by 45% compared to the parental DMS3 phage at 8 h post-infection (hpi) (Fig. 1 and S1). Moreover, compared with the parental DMS3 phage, both DMS3 AqdC alone and the combined anti-QS phage treatment significantly reduced PQS accumulation, by 87-90% and 70-76%, respectively, at 6-10 hpi, as quantified using the PqsR-responsive bioreporter (Fig. 1 and S1). Overall, although the engineered anti-QS phages did not alter 3O-C12-HSL levels under the experimental conditions, the combined DMS3 AqdC and DMS3 QsdA treatment significantly suppressed both C4-HSL and PQS accumulation, highlighting its strong capacity to disrupt QS signaling during infection.

Phage-resistant mutants are naturally selected for during phage exposure and represent a significant challenge in phage therapy [10]. Therefore, we challenged the robustness of our anti-QS phages to inhibit QS activation in mixed populations of phage-susceptible PA14 and DMS3 phage-resistant PA14 cells. As phage receptors perform key functions in bacteria, including virulence [39], we created a phage-resistant mutant without deleting genes required for the production of the phage receptor, type IV pili [40], to avoid adverse effects. Instead, we relied on a DMS3 prophage incapable of producing phage particles to provide phage resistance and additionally mutated the *aqs1* gene to prevent the prophage from inhibiting LasR. Specifically, we deleted the DMS3 genes *gp26-48* in a PA14 DMS3 *aqs1*_FSDARE lysogen. The PA14 DMS3 *aqs1*_FSDARE Δ*gp26-48* strain (PA14^R^ hereafter) was resistant to DMS3 phage-mediated killing, unlike the PA14 WT strain, as expected (Fig. S2).

We then tested the ability of the DMS3 anti-QS phages to effectively silence mixed populations consisting of 90% phage-susceptible PA14 WT and 10% phage-resistant PA14^R^ (Fig. 1). As in the PA14 monoculture, anti-QS phages lysed the 10% resistant co-cultures to a similar extent as the parental DMS3 phage (Fig. 1). Consistent with the PA14 monoculture results, anti-QS phage infection did not significantly reduce 3OC12-HSL levels in 10% resistant co-cultures compared with the parental DMS3 phage under the experimental conditions (Fig. 1 and S1). Yet co-infection with DMS3 QsdA and AqdC significantly reduced C4-HSL levels by 35% in 10% resistant co-cultures at 6 hpi compared to DMS3 parent phage. Correspondingly, infections of the 10% resistant co-cultures with either DMS3 AqdC alone or the combination of DMS3 AqdC and QsdA, significantly decreased PQS accumulation by 37-63% at 6-10 hpi and 34-40% at 8-10 hpi, respectively, compared to the DMS3 parental phage (Fig. 1 and S1). Although they did not significantly reduce 3OC12-HSL levels, the combined DMS3 QsdA and DMS3 AqdC treatment lowered C4-HSL accumulation, and both DMS3 AqdC alone and the dual phage treatment effectively suppressed PQS levels. Overall, QS inhibition in the 10% resistant co-cultures followed the same qualitative pattern observed in phage-susceptible monocultures, albeit with generally reduced magnitude. Notably, DMS3 AqdC consistently produced the strongest PQS inhibition across both culture types, whereas significant repression of C4-HSL required the combined activity of QsdA and AqdC. Collectively, these results demonstrate that the anti-QS phages degrade QS molecules in phage-resistant subpopulations through infection of phage-susceptible cells, enabling production of quorum quenching enzymes, reducing QS signals in the environment.

### Prophage-encoded QsdA and AqdC reduce QS-regulated virulence factor production

Having established that our anti-QS phages disrupt QS molecule accumulation, we next examined the extent to which they modulate downstream QS-regulated virulence factor production. Specifically, we quantified elastase activity and pyocyanin production, activated by LasR and PqsR respectively, in PA14 DMS3 lysogens in which the Aca repressor was removed. The strains included PA14 DMS3 *aqs1_FSDARE* (Parent lysogen), PA14 DMS3 *aqs1_FSDARE ΔacrIE-aca::qsdA* (QsdA lysogen) and PA14 DMS3 *aqs1_FSDARE* Δ*acrIE-aca::aqdC* (AqdC lysogen) (Fig. 2). We quantified elastase activity of sterile lysogen supernatants using an elastin-Congo red assay. The DMS3 AqdC lysogen exhibited elastase activity comparable to the parent lysogen, indicating that expression of *aqdC* did not alter elastase activity (Fig. 2A). In contrast, elastase activity was significantly decreased by 31% in the DMS3 QsdA lysogen compared to the parent lysogen (Fig. 2A), demonstrating that expression of *qsdA* from the DMS3 prophage attenuates LasR-regulated virulence factor production, consistent with elastase being primarily controlled by the LasR system. We next quantified the PQS-regulated virulence factor pyocyanin in the lysogens. Pyocyanin levels in the DMS3 QsdA lysogen were comparable to those of the parent lysogen, whereas the DMS3 AqdC lysogen showed a significant 43-44% reduction in pyocyanin production (Fig. 2B). This demonstrates that expression of *aqdC* from the DMS3 prophage attenuates PQS-regulated virulence factor production, which is in line with pyocyanin production being activated by PqsR. Together, these results show that QsdA and AqdC each selectively disrupt distinct QS-regulated virulence outputs when expressed from the DMS3 prophage, highlighting the potential of the anti-QS phages in reducing virulence factor production.

**Figure 2.**
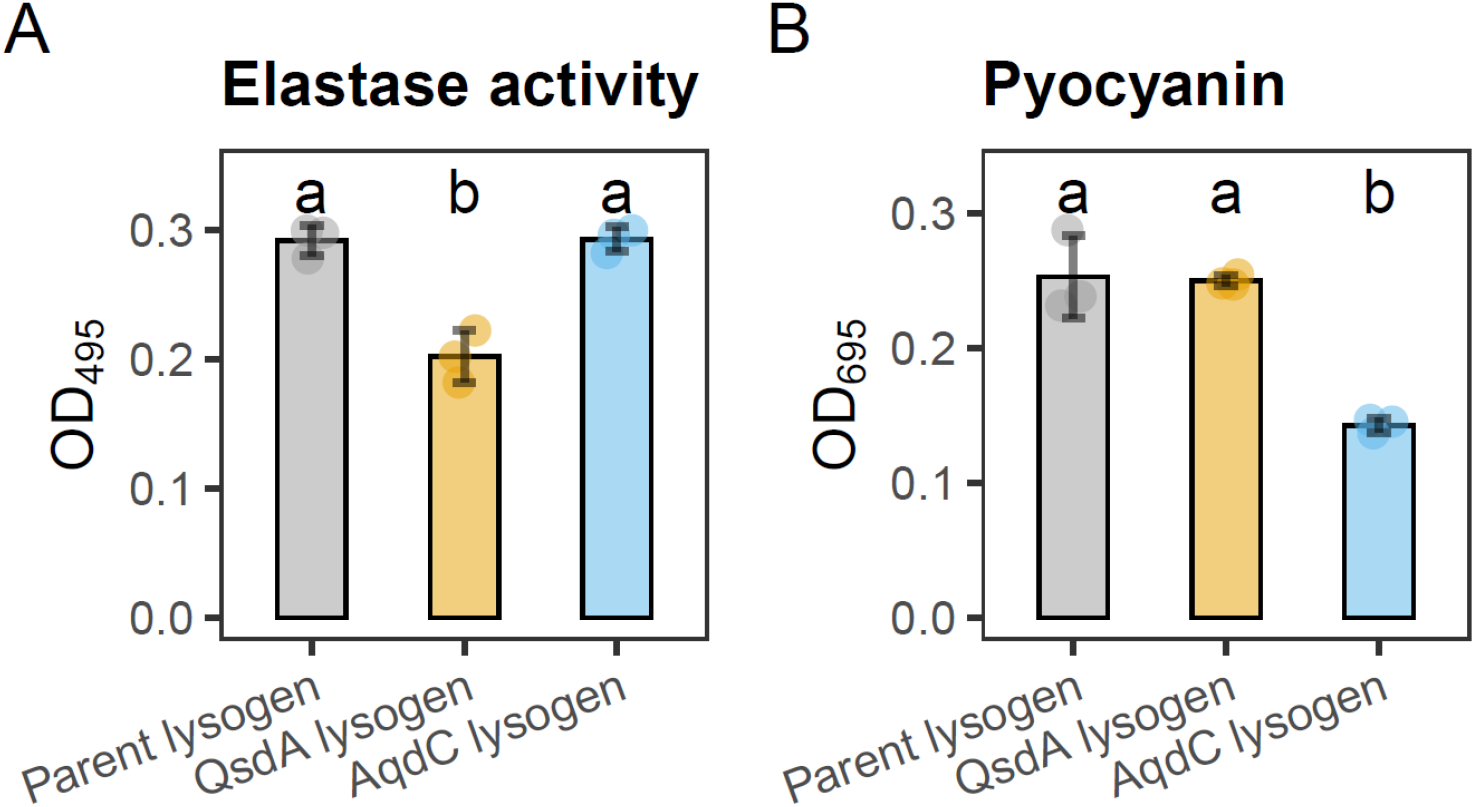
Virulence factor production is reduced in anti-QS PA14 lysogens. Elastase activity **(A)** and pyocyanin **(B)** were quantified from culture supernatants of lysogens carrying DMS3 parent- or engineered anti-QS prophages expressing *qsdA* or *aqdC*. Elastase activity was quantified using an elastin-Congo red assay measuring absorbance at 495 nm, and pyocyanin levels were determined by measuring absorbance at 695 nm. Error bars represent standard deviation from biological replicates (*n*=3). One-way ANOVA was used to test for overall differences among strains, followed by Tukey’s HSD test to obtain all pairwise comparisons. Groups not significantly different (α=0.05) share the same letter.

### Anti-QS phages enhance survival of *P. aeruginosa*-infected *G. mellonella*

After confirming the ability of anti-QS phages and lysogens, respectively, to inhibit QS and suppress virulence factor production *in vitro*, we assessed whether these effects persisted *in vivo*, considering that infection dynamics are shaped by spatial structure and complex tripartite interactions between phages, bacteria, and eukaryotic hosts which can lead to outcomes distinct from those observed in simplified laboratory systems. Here, we used a *G. mellonella* wax moth larvae burn wound model of *P. aeruginosa* infection, since it closely mimics the infection dynamics of mammalian infection models [41]. To investigate the ability of our anti-QS phages to rescue *G. mellonella* from *P. aeruginosa* infection, we either control treated burn wounds or infected them with PA14. One hour after initiating infection, some infected larvae received treatment either with the parental DMS3 or with engineered anti-QS phages (Fig. 2). We monitored larval survival over time. Phage only controls were indistinguishable from untreated and burn wound only controls, indicating that neither the burn procedure nor phage application alone affected larval viability (Fig. 2A). On average, 91% of larvae across all control treatments survived the course of the 96-h experiment. In contrast, PA14 infection alone significantly reduced survival to 37%. When treating larvae with the DMS3 parent phage, 34% of larvae survived PA14 infection, indicating that the parental phage did not rescue larvae from PA14 killing. Strikingly, treatment with the anti-QS phages DMS3 QsdA and AqdC increased survival to 67% - a 100% relative improvement over DMS3 parent treatment. Because survival outcomes with the individual anti-QS phages DMS3 QsdA or AqdC alone were not significantly different from their combined treatment, subsequent experiments were conducted using a 1:1 mixture of both anti-QS phages. To further challenge the robustness of our phages, we infected *G. mellonella* larvae with PA14 co-cultures containing 10% phage-resistant PA14^R^ cells (Fig. 2B). To account for batch variability, larvae were again control-treated with a mixture of all DMS3 phages in the absence of infection, resulting in 87% of larvae remaining viable over the course of the 96-h incubation. Infection with PA14 co-cultures reduced larval survival to 20%. While 27% of larvae survived infection when treated with the DMS3 parent phage, a non-significant improvement of survival, treatment with anti-QS phages increased survival to 54% - a 100% relative improvement over DMS3 parent treatment. These results collectively highlight the therapeutic potential of engineered anti-QS phages in attenuating *P. aeruginosa* PA14 virulence and enhancing survival *in vivo*, even in the presence of phage-resistant subpopulations.

Taken together, our results show that engineered anti-QS phages can reduce the accumulation of QS signals (Fig. 1), suppress the production of key virulence factors (Fig. 2), and substantially improve survival of PA14-infected *G. mellonella* (Fig. 3). By targeting both AHL- and PQS-dependent QS pathways, QsdA- and AqdC-expressing phages effectively weaken the ability of *P. aeruginosa* to coordinate pathogenic behaviors. These findings establish engineered anti-QS phages as a powerful strategy for disrupting bacterial communication and attenuating virulence in complex infection environments.

**Figure 3.**
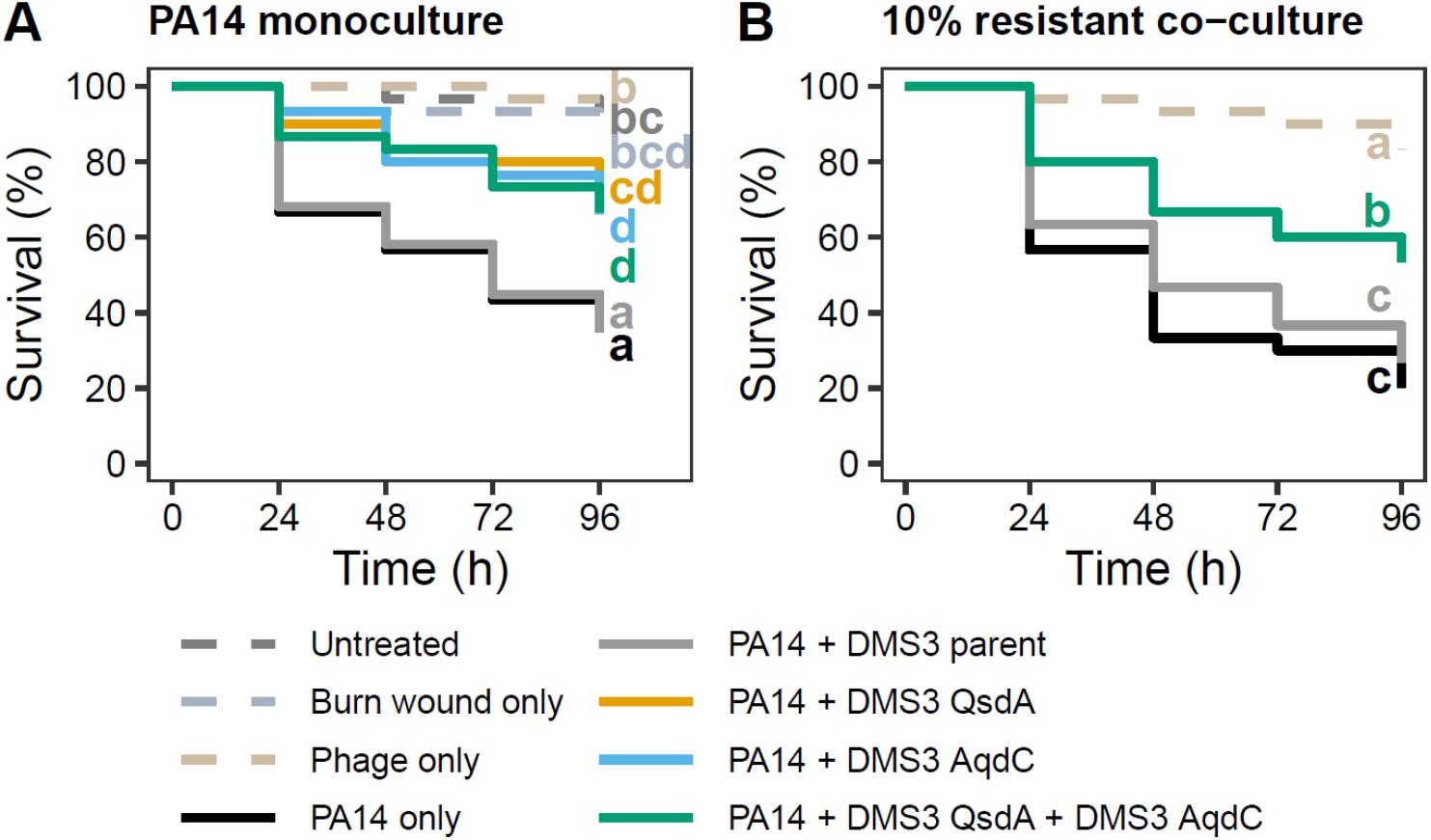
Virulent anti-QS phages protect *G. mellonella* larvae from *P. aeruginosa* killing. **(A)** The effect of our anti-QS phages was assessed using a *G. mellonella* larvae burn infection model. Larvae were either untreated or inflicted with a burn wound. Bun wounds were either left untreated, treated with phage only, or infected with PA14. The PA14-infected wounds were untreated or treated 1 h post-infection at an MOI of 100 with either the DMS3 parent, DMS3 QsdA, DMS3 AqdC, or a 1:1 mixture of the engineered anti-QS phages. **(B)** As in A, but wounds were either treated with phage only or infected with PA14 co-cultures containing 10% phage-resistant PA14^R^ cells, followed by treatment with the DMS3 parent or a 1:1 mixture of the engineered anti-QS phages. Statistical significance was determined using Mantel-Cox log-rank tests. *n*=30 larvae per condition.

## Discussion

The global threat of multidrug-resistant bacteria urgently demands new antimicrobials [1, 42]. Phage therapy is one option [5, 6] however, bacteria fight back, which may reduce the effectiveness of phage therapy. Here we present engineered anti-QS phages as precision antimicrobials that combine the unconventional treatment options of quorum quenching and phage therapy to treat *P. aeruginosa* infections in *G. mellonella* wax moth larvae (Fig. 3). This synergistic combination allows specific pathogen killing alongside production and release of the anti-QS enzymes QsdA and AqdC at the site of infection. This approach may enhance the antimicrobial effect of phages by silencing the bacteria evolving immunity to the phages, as QS molecule-degrading enzymes released from infected phage-sensitive bacteria degrade the extracellular QS molecules emitted by any phage-immune bacteria, hindering activation of virulence of the collective *P. aeruginosa* population (Fig. 2). In a patient setting, this may pave the way for natural clearance of residual phage-resistant mutants by the human immune system and/or healthy microbiota.

An early study combined phage therapy and quorum quenching by equipping the coliphage T7 with the AiiA lactonase to infect *Escherichia coli*, which does not itself produce homoserine lactones. The T7 phage inhibited biofilm formation of an *E. coli-P. aeruginosa* mixed species biofilm [43]. Here, we show that the anti-QS enzymes QsdA and AqdC expressed from virulent *P. aeruginosa* phages effectively limit elastase and pyocyanin virulence factor production (Fig. 2) and disarm *P. aeruginosa* when infecting *G. mellonella* larvae (Fig. 3). While previous studies have shown that expressing quorum quenching enzymes from plasmids limits *P. aeruginosa* virulence effectively, applying purified quorum quenching enzymes to treat *P. aeruginosa* infections of various animal models have shown limited success, in part due to challenges with delivery [34, 44–46]. This limited success could additionally be due to the anti-QS enzymes being degraded or extensively diluted in the animal model before reaching the site of infection. Our approach circumvents these limitations by enabling *in situ* amplification of the “drug” at the site of infection, using a phage specific for the pathogen, in addition to providing phage-mediated clearance of the targeted pathogen. A key advantage of using quorum-quenching enzymes such as QsdA and AqdC, lies in their ability to act beyond the boundaries of the infected, phage-susceptible cell. Intracellular QS inhibitors degrade or block QS molecules only inside the specific phage-infected bacterial cell. Consequently, their activity is inherently restricted to the fraction of the population that is phage-sensitive and infected at the time. In contrast, the enzymes delivered by our engineered phages degrade signal molecules released by both phage-susceptible and phage-resistant subpopulations. This extracellular mode of action enables suppression of QS-regulated virulence at the population level, even among bacteria avoiding phage infection. Thus, the anti-QS enzymes effectively bridge the gap between infected and uninfected bacteria, creating a community-wide anti-virulence effect that purely intracellular QS inhibition strategies cannot achieve.

As QS controls virulence properties in a broad range of pathogens infecting humans, animals, and plants [47], the concept of combining quorum-modulating proteins [48] and phage therapy may be generalized to target other bacterial pathogens beyond the human pathogen *P. aeruginosa* studied here. The principle of equipping phages with the ability to manipulate their host bacteria for reduced bacterial virulence and enhanced bacterial killing may enhance the overall effectiveness of future phage therapy efforts.

## Acknowledgements

N.M.H-K. was supported by the Independent Research Fund Denmark grant 1054-00099B and N.M.H-K and J.J.M. were supported by the Novo Nordic Foundation grant NNF24OC0096270.

## Methods

### Bacterial growth conditions

*P. aeruginosa* and *E. coli* strains were grown at 37 °C with 300 rpm shaking in LB (10 g L^-1^ tryptone, 5 g L^-1^ yeast extract, and 10 g L^-1^ NaCl) or on LB agar plates containing 1.5% agar (wt/vol) unless otherwise specified. When appropriate, overnight cultures were diluted in fresh LB to a lower optical density (OD600). Bacterial strains, plasmids, and phages used are listed in Table 1.

### DNA manipulations

Genes encoding the N-acyl-homoserine lactonase QsdA from *Rhodococcus erythropolis* (accession no. AP008957) and the PQS dioxygenase AqdC from *Mycobacteroides abscessus* (accession no. CP050978) were synthesized by TAG Copenhagen or Eurofins, respectively. Constructs were inserted downstream of the anti-CRISPR promoter in the DMS3 phage, while deleting the type I-E anti-CRISPR gene (DMS3-30) but leaving the anti-CRISPR-associated repressor gene (DMS3-31) intact in lysogens carrying DMS3 prophages. Subsequently, the phages were made virulent by deleting bp position 154-380 in the *cI* gene (DMS3-1) [36] as detailed below. For PA14 DMS3 lysogens expressing *qsdA* or *aqdC*, the anti-CRISPR-associated repressor gene (DMS3-31) was removed.

Plasmids were extracted using NucleoSpin® Plasmid EasyPure kit (Machery-Nagel, Ref. 740727.250). Plasmids for cloning were digested with FastDigest™ restriction enzymes (Thermo Scientific™) and dephosphorylated using FastAP Thermosensitive Alkaline Phosphatase (Thermo Scientific™, Ref. EF0654) according to manufacturer’s instructions. Linearized plasmids were gel-purified using NucleoSpin® Gel and PCR Clean-up kit (Machery-Nagel, Ref. 740609.250). For cloning purposes, PCRs were carried out using the Q5® High Fidelity DNA Polymerase (New England BioLabs, Ref. M0491L). For other purposes, PCRs were performed using the DreamTaq DNA Polymerase (Thermo Scientific™, Ref. EP0702). PCR products were purified using NucleoSpin® Gel and PCR Clean-up kit (Machery-Nagel, Ref. 740609.250). Desired inserts were assembled with digested plasmids using T4 DNA Ligase (Thermo Scientific™, Ref. EL0016) or by Gibson assembly with fragments containing 50 bp overlaps, using the NEBuilder® HiFi DNA Assembly kit (New England BioLabs, Ref. M5520AA), following the manufacturer’s instructions. Plasmids were recovered in One Shot™ TOP10 Chemically Competent Cells (Invitrogen, Ref. C404003) according to manufacturer’s instructions or in *E. coli* SM10 λ*pir* using TSS chemical competence[49]. Plasmids were confirmed by PCR and sequencing (Eurofins and/or Plasmidsaurus).

The pEXG2-based plasmids were conjugated into *P. aeruginosa* strains by mating. Exconjugants were selected on LB containing 30 µg mL^-1^ gentamicin and 100 µg mL^-1^ irgasan and cultured at 37 °C overnight. Then, colonies were grown in LB at 37 °C for 2 h, plated on LB containing 15% sucrose and no NaCl and cultured at 37 °C overnight. Mutants were confirmed by PCR and sequencing (Eurofins).

To insert gRNA into pAB01, primer pairs were phosphorylated and annealed in 50 mM NaCl at 98 °C for 5 min, followed by gradually cooling down 2 °C for 30 s at each temperature. The annealed primers were then ligated into BbsI-digested and dephosphorylated pAB01 using T4 ligase. Subsequently, homology templates for deleting *cI* (DMS3-1) were fused using PCR and inserted into NheI-digested pAB01 using Gibson assembly. The pAB01-based plasmids were electroporated into *P. aeruginosa* PA14 pre-washed in 300 mM sucrose and recovered in LB at 37 °C for 1 h and then plated on LB containing 50 µg mL^-1^ gentamicin.

Virulent DMS3 phages were constructed by infecting 1:1,000 back-diluted overnight cultures of *P. aeruginosa* PA14 pAB01-DMS3-Δ*cI* with 200 µL of sterile-filtered supernatants from overnight cultures of various PA14 DMS3 lysogens. Infections were caried out in 2 mL LB containing 50 µg mL^-1^ gentamicin and 0.1% arabinose at 37 °C for 5 h. To isolate single plaques, plate lysates were created using 500 µL supernatant and 200 µL uninfected *P. aeruginosa* PA14 pAB01-DMS3-Δ*cI* overnight culture in soft agar containing 50 µg mL^-1^ gentamicin and 0.1% arabinose and incubated at 37 °C overnight. Virulent DMS3 phages were confirmed by PCR, and sequencing, and individual plaques were re-streaked on lawns of PA14 pAB01-DMS3-Δ*cI*.

### Phage proliferation

Briefly, the engineered phages were purified by re-streaking single plaques on PA14 lawns three times, resuspended in SM buffer (100 mM NaCl, 8 mM MgSO4, 50 mM Tris HCl, pH 7.5, 0.01% gelatin), and proliferated using plate lysates with PA14 Δpf5 Δ*lasI* Δ*rhlI* Δ*pqsA* lawns, to obtain QS-free lysates [38]. In more detail, the single plaques were resuspended in 500 µL SM buffer and after three rounds of re-streaking, 300 µL of the resuspended plaque solution was mixed with 50 µL PA14 overnight culture and 5 mL molten soft agar, subsequently poured onto LB agar plates and incubated at 37 °C overnight. Next, 5 mL SM buffer was added to the infected lawns and incubated for 3-5 h at room temperature. The buffer was then collected, 0.2-µm sterile-filtered and phages were enumerated using standard plaque assay on lawns of PA14 or PA14 pAB01-DMS3-Δ*cI* containing 50 and 200 uL overnight culture, respectively. For lawns of PA14 pAB01-DMS3-Δ*cI*, the soft agar was supplemented with 50 µg mL^-1^ gentamicin and 0.1% arabinose.

### Infection assays

To assess the phage resistance conferred by the modified DMS3 prophage in PA14^R^, overnight cultures of *P. aeruginosa* PA14 and PA14^R^ were grown in LB at 37 °C with shaking at 300 rpm. Cultures were back-diluted to OD_600_ of 0.01 and either left uninfected or challenged with the DMS3 parent phage at an MOI of 10 (1*10^8^ PFU ml^-1^). Cultures were incubated at 37 °C with intermittent shaking. Absorbance measurements were recorded using a Synergy H1 plate reader (BioTek Instruments Inc.) with software Gen5 v3.05.11.

To obtain culture supernatants for QS signal measurements, *P. aeruginosa* PA14 and PA14^R^ overnight cultures prepared in LB containing 10 µM ZnSO_4_ were back-diluted to OD_600_ of 0.001 in LB with 10 µM ZnSO_4_ and grown at 37 °C shaking with 300 rpm. For the co-culture assay, PA14 and PA14^R^ were mixed in a 9-to-1 ratio based on OD_600_. At OD600 ~ 0.2, the cultures were challenged with a total MOI of 10 (2*10^9^ PFU ml^-1^) with either DMS3 parent or variants expressing either *qsdA, aqdC* or a mixture of *qsdA* and *aqdC*-encoding variants. After OD_600_ monitoring, sterile-filtered supernatants were stored at –20 °C for up to 2 days and subsequently measured using the QS bioreporter assay.

### Quorum sensing luminescence reporter assays

To evaluate the relative abundances of the QS molecules 3OC12-HSL, C4-HSL, and PQS in sterile-filtered culture supernatants, we used *E. coli* reporter strains carrying an arabinose-inducible pBAD vector and a pCS26 *lux* reporter. In these strains, arabinose induces transcription of genes encoding either the 3O-C12-HSL receptor LasR, C4-HSL receptor RhlR, or PQS receptor PqsR, promoting transcription of the *lasB, rhlA*, or the *pqsA* promoters, respectively, upstream of the *luxCDABE* operon in pCS26 [37, 38]. Thus, these strains luminesce in response to the respective QS molecules. Briefly, overnight cultures of the *E. coli* strains carrying the respective plasmids were diluted 100-fold in LB containing 100 µg mL^-1^ kanamycin, 100 µg mL^-1^ ampicillin, and 0.1% arabinose. To each well, 10 µL *P. aeruginosa* sterile-filtered culture supernatant was added, resulting in a total volume of 200 µL per well. For PQS measurements, the *P. aeruginosa* supernatants were diluted 100-fold, whereas for measurements of 3O-C12-HSL and C4-HSL, the supernatants were undiluted. The measurements were carried out in black-well, clear-bottom 96-well plates with intermittent shaking at 30 °C. OD_600_ and luminescence was measured using a Synergy H1 plate reader (BioTek Instruments Inc.) with software Gen5 v.3.05.11.

Luminescence values were first normalized by dividing by the corresponding OD_600_ measurements. For each QS signal, a fixed time window was defined based on the earliest and latest peak times observed across all treatments. The average luminescence/OD_600_ value across this window was then calculated for each sample. Specifically, the averaging windows were 7–22 h for 3OC12-HSL reporter, 5–24 h for C4-HSL reporter, and 2.67–7 h for PQS reporter. These representative values were then normalized within each experiment to the DMS3 parent treatment: to the 0 hpi value for 3OC12-HSL and PQS, and to the 2 hpi value for C4-HSL due to zero values at 0 hpi.

Time series data of relative QS signal values from the QS bioreporter assays were analyzed using generalized additive mixed models (GAMMs) implemented in mgcv [50]. For each culture condition and QS signal, the model included phage treatment as a fixed effect, treatment-specific smooth functions of hours post-infection, and biological replicate as a random effect. Pairwise differences between treatments were estimated from the fitted model using the linear predictor matrix, which propagates uncertainty from all model coefficients. Contrasts were evaluated over a dense grid of timepoints, and 95% confidence intervals were derived from the model covariance matrix. Time intervals for which the confidence interval did not include zero were interpreted as significant differences. For visualization, estimated differences and confidence intervals were plotted for each QS reporter value and culture type, with the dashed zero line representing no difference. All analyses and data processing were performed in R using mgcv v. 1.8-38, [51] ggplot2 v. 3.5.2, and tidyverse v. 2.0.0 [52].

### Elastase virulence assay

Elastase virulence factor production was determined in PA14 DMS3 lysogens (parent, QsdA, AqdC). The protocol was adapted from [53] with modifications. Cultures were grown overnight in LB to an OD_600_ of 3.0 and sterile filtered using 0.2 µm filters. 50 µL of supernatant was added to tubes containing 950 µL elastase buffer (100 mM Tris-HCl, 1 mM CaCl_2_, pH 7.5) and 2.5 mg elastin-Congo red (Sigma-Aldrich, E0502). After incubation in the dark at 37 °C with shaking at 300 rpm for 2 h, samples were centrifuged for 5 min. Elastase activity in supernatants was quantified by measuring absorbance at 495 nm using a spectrophotometer. Measurements were performed in biological triplicates.

### Pyocyanin virulence assay

Selected strains were cultured in LB at 37 °C with shaking at 300 rpm for 18 h. Cells were pelleted by centrifugation at 13,000 rpm for 3 min, and OD_695_ of the supernatant was measured as a proxy for pyocyanin, a green pigment that can be quantified by absorbance at this wavelength. Absorbance measurements were obtained using a Synergy H1 plate reader (BioTek Instruments Inc.) with software Gen5 v3.05.11.

### Larvae burn wound infection assay

*Galleria mellonella*, greater wax moth larvae, reared without antibiotics, were obtained from a local pet supplier. Upon arrival, larvae were stored at 12 °C for no longer than 10 days. Prior to experiments, larvae were size-matched (2.5-3.0 cm in length, respectively) and examined visually; only larvae showing signs of movement and no melanization were used. Selected larvae were placed in sterile Petri dishes (two larvae per dish) and surface-sterilized with 70% ethanol. To facilitate handling, larvae were briefly chilled at 4 °C for a maximum of 10 min to restrict movement. Burn wounds were generated as previously described, adapted from Maslova et al., 2020. Briefly, the head of a stainless-steel nail (2 mm diameter) was heated to red hot using a Bunsen burner and allowed to cool for 4 s before gentle application to the dorsal surface of each larva for 4 s. Fresh wounds were immediately infected with 10 µL of a standardized PA14 inoculum (OD_600_ of 1.0), which was slowly pipetted directly onto the wound. Following infection, larvae were incubated at 37 °C to allow establishment of infection. 1 h post-infection, selected larvae received phage treatment at an MOI of 100. Control groups included untreated larvae, larvae with uninfected burn wounds, and larvae receiving phage treatment in the absence of bacterial infection. Larvae were kept at 37 °C for 4 days and monitored for survival, based on movement and melanization, at the indicated time points.

## Figures and Figure legends

**Figure S1.**
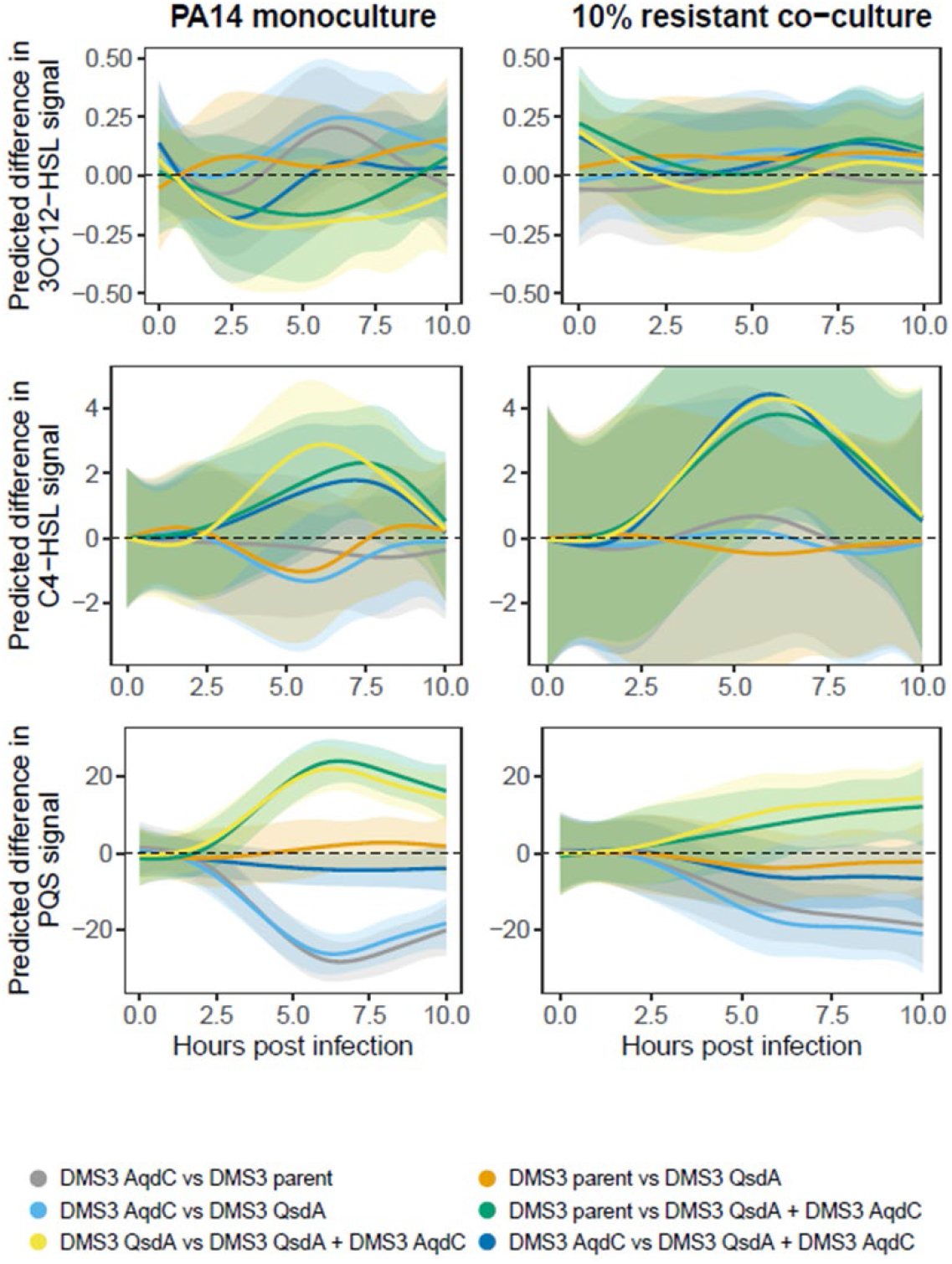
GAMM-based pairwise differences in QS bioreporter activity. Generalized additive mixed models (GAMMs) were fitted to assess how QS reporter outputs differed among phage treatments through time. For each reporter and input culture type, pairwise treatment differences are shown along with 95% confidence intervals calculated from the model’s full covariance matrix. The dashed horizontal line marks zero difference. Periods where the confidence interval does not include zero indicate statistically significant differences between treatments. All modelling and figure generation were conducted in R as described in the Methods.

**Figure S2.**
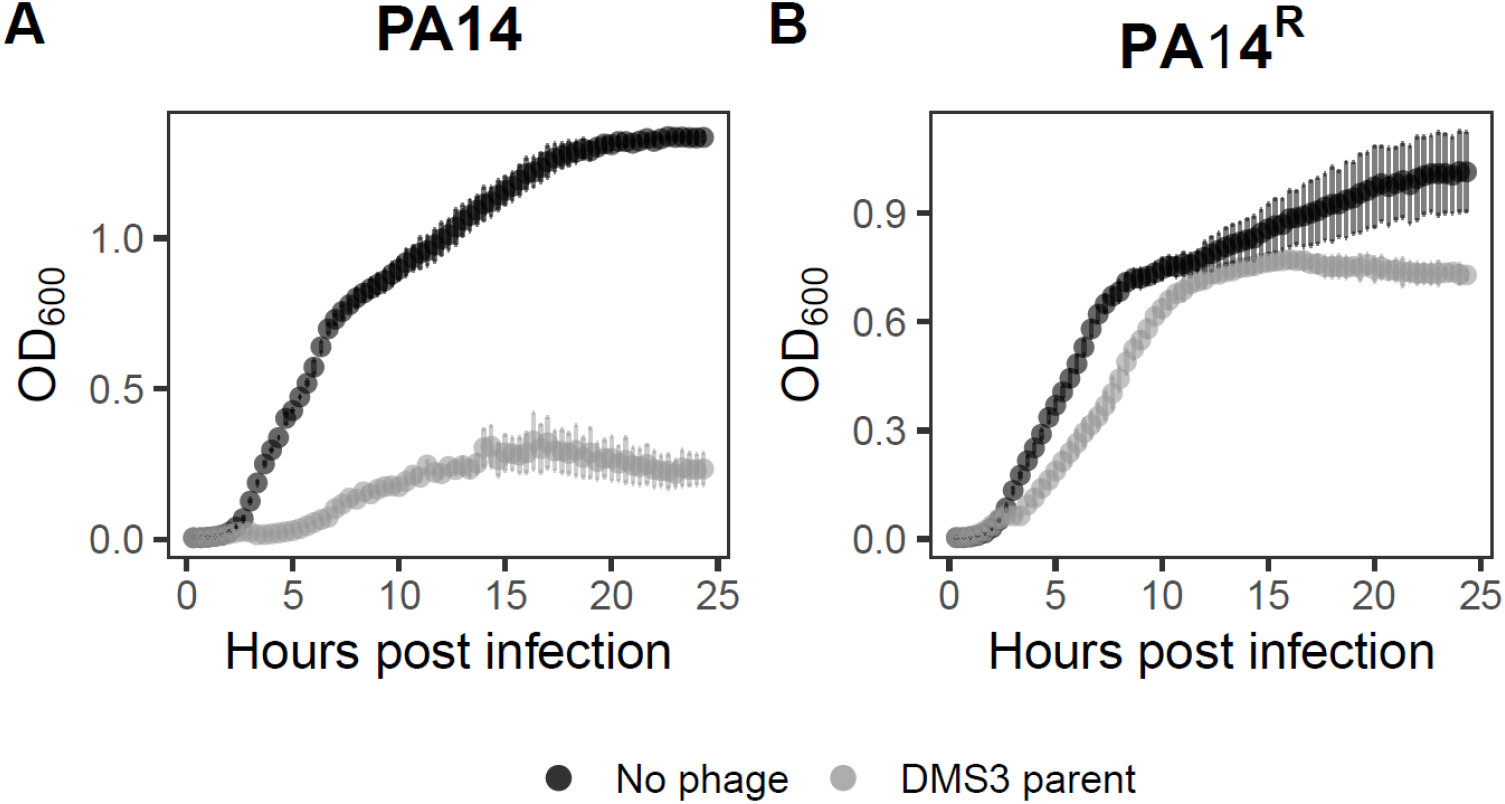
*P. aeruginosa* PA14^R^ is resistant to DMS3 parent killing. Overnight cultures of *P. aeruginosa* PA14 and PA14^R^ were diluted to an OD_600_ of 0.01, and either left uninfected or challenged with parental DMS3 phage (MOI=10; 1*10^8^ PFU ml^-1^). Bacterial abundance was monitored by OD_600_. Error bars represent standard deviation from biological replicates (*n*=3).

**Table S1.**
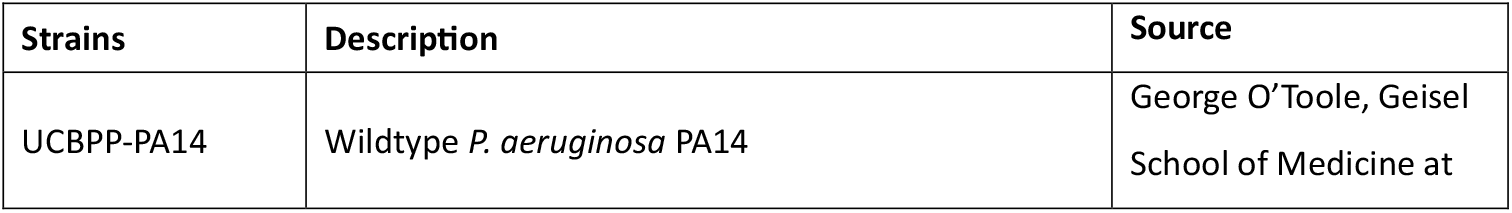
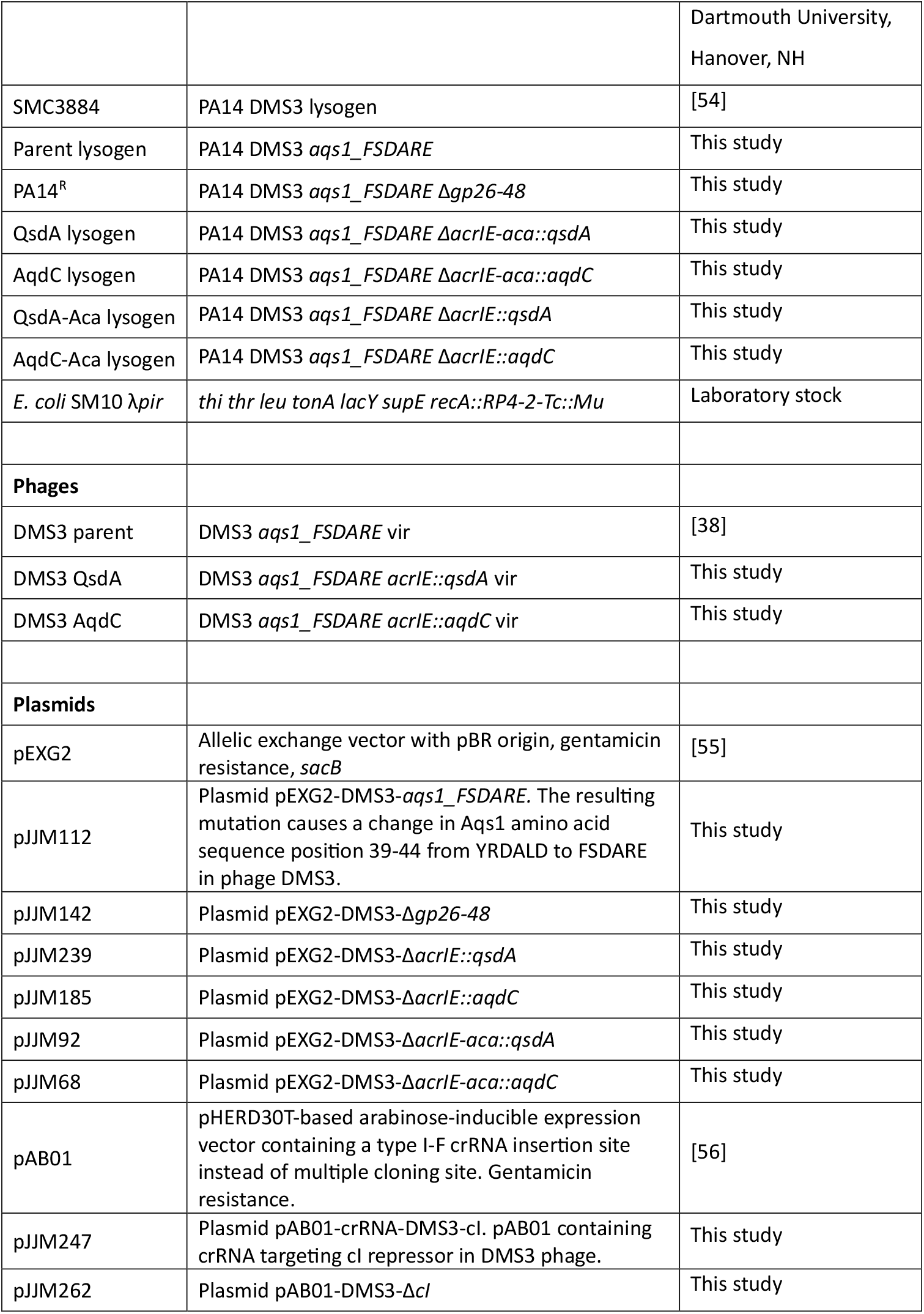
Strains, phages, and plasmids used in this study.

